# Mechanisms of fork stalling and reversal at pathogenic CANVAS (AAGGG)_n_ repeats

**DOI:** 10.64898/2026.10.27.762263

**Authors:** Nicholas H. Mandel, Sofia Movafaghi, Rishika Vaid, Olga Usalka, Simone Ummarino, Joe M. Polyak, Masayoshi Honda, Negin Mozafari, Rula Zain, Maria Spies, Sergei M. Mirkin

## Abstract

Expansions of the pathogenic (AAGGG)n repeat within the RFC1 gene cause the hereditary disease CANVAS. We have shown that this repeat impedes DNA replication both *in vitro* and *in vivo*. Here, we investigated the mechanisms of replication fork stalling and reversal at (AAGGG)n repeats within an SV40-based episome in human cells. A comprehensive 2dimensional electrophoretic analysis of replication intermediates revealed a strict orientation dependence of the replication block, evident only when the homopurine sequence was in the lagging-strand template. Fork accumulation was further increased in the presence of the G4DNA stabilizer pyridostatin, implicating G4-DNA in the process. This stalling was followed by an accumulation of reversed forks and further analysis revealed roles for SMARCAL1 and RAD51 in this process. Biochemical studies showed that SMARCAL1, with assistance from RPA, efficiently remodels a fork carrying the G4-forming AAGGG repeat in the ssDNA gap, but not a substrate carrying the repeat in triplex conformation. We conclude that stalling results from G4-DNA formation by the repeat, which then triggers SMARCAL1-mediated fork reversal at the repeat. Given the mirror symmetry of the sequence, the nascent strands at the reversed forks likely adopt a triplex conformation, leading to a thorough accumulation of branched structures.

## Introduction

Microsatellites or short tandem repeats (STRs) constitute up to 7% of the human genome^1,2^ and play important roles in gene expression^3,4^. At the same time, STRs often have higher mutation rates than non-repetitive DNA^5^, resulting in repeat instability, *i.e.* their expansions or contractions^6^. Large-scale repeat expansions have been linked to more than 60 genetic disorders^7–11^, cumulatively called repeat expansion disorders (REDs).

STRs can form different alternative DNA structures^12–14^ depending on their sequence composition. These include quadruplex G4-DNA formed by some G-rich STRs^15,16^, triplex HDNA formed by homopurine-homopyrimidine repeats^17–19^, DNA cruciforms primarily formed by (AT_)n_ repeats^20–24^, and slipped-strand S-DNA formed by (CNG_)n_ repeats^25^. Given that DNA primarily exists in its double-stranded B-form, duplex unwinding is a prerequisite for the formation of all alternative DNA structures^26^. Therefore, they are dynamic structures that can be transiently formed throughout the course of major genetic transactions including DNA replication, transcription, repair, and recombination^27–29^. During DNA replication, formation of alternative DNA structures may create impediments to the replisome, resulting in replication fork stalling and reversal^30–32^. In the event the replisome has not disassembled, the latter provides a convenient opportunity for fork restart; however stronger stalling resulting in fork collapse may require more mutagenic pathways like break-induced replication^33^. It is generally believed, therefore, that formation and resolution of dynamic DNA structures formed by pathogenic STRs is at the heart of REDs.

While most REDs are caused by trinucleotide repeat expansions, longer expandable repeats have attracted significant attention within the last decade^34^. One recent example is the biallelic expansion of the (AAGGG)_n_ repeat within the second intron of the *RFC1* gene, causing a neurological disease called CANVAS: **c**erebellar **a**taxia with **n**europathy and **v**estibular **a**reflexia **s**yndrome^35,36^. It is a rare, late-onset autosomal recessive disorder that often presents with symptoms including difficulty walking, chronic cough, oscillopsia, and autonomic dysfunction in older adults^37^. CANVAS differs from other REDs in that unaffected individuals contain the nonpathogenic (AAAAG)_n_ repeat, which has a limited potential to expand, while affected individuals contain massively expanded (ACAGG)_n_, (AAAGG)_n_, or (AAGGG)_n_ repeats within the *RFC1* gene^38–40^. Notably, biallelic expansions of (AAGGG)_n_ repeats ranging from 200 to over 2,000 units account for 92% of CANVAS cases^35^. Recently, large biallelic (AAGGG)_n_ repeat expansions in the *RFC1* gene have also been identified in Parkinson’s disease patients^41^.

Thorough cohort studies showed that the frequency of carriers having monoallelic (AAGGG)_n_ repeat expansions in the *RFC1* gene reaches 4% in European populations^42^. These carriers are asymptomatic, suggesting a recessive inheritance trait, which is usually due to a loss-of-function mutation. Yet, initial studies of CANVAS patients do not show any significant decrease in *RFC1* expression35. Studies in patient-derived neurons point to repeat-dependent and *RFC1*independent synaptic dysregulation^43^, but the mechanism remains unknown.

Investigation into whether cells can faithfully duplicate long repetitive (AAGGG)_n_ sequences, or fail to do so, could help unravel the mechanisms of repeat expansion leading to CANVAS. Since AAGGG is a homopurine-homopyrimidine mirror repeat, which also contains regularly spaced G_3_ runs, it has the propensity to form either H-DNA or G4-DNA structures. Circular dichroism spectroscopy has indeed confirmed formation of both structures specifically by the pathogenic repeat, but not by its nonpathogenic counterpart^44^. Our lab has previously investigated replication through (AAGGG)_n_ repeats both *in vitro* and in *vivo*^45^. We found that even short (AAGGG)_10_ repeats in the template DNA strand stalled DNA polymerization *in vitro*. The fine pattern of the polymerization stalling was consistent with G4-DNA formation in the presence of K^+^ ions or HDNA formation in the presence of Mg^2+^ ions. (CCCTT)_n_ runs in the template DNA strand, or non-pathogenic DNA repeats didn’t cause polymerization blockage *in vitro*. Using 2dimensional gel-electrophoresis (2DGE) to analyze replication intermediates, we found that pathogenic repeats caused strong replication stalling in an SV40-derived episome in human cells. Strikingly, we also observed a profusion of branched replication intermediates of the same size as stalled intermediates, which are generally recognized as reversed replication forks^46,47^.

The sheer magnitude of the (AAGGG)n repeat-mediated replication hindrance prompted us to investigate its genetic controls and underlying mechanisms. Our comprehensive 2DGE analysis revealed a length- and orientation-dependent replication blockage, evident only when a repeat reached its carrier size and its homopurine run was in the lagging-strand DNA template. Fork stalling appears to depend on G4-DNA formation, as it becomes even stronger in the presence of the G4-DNA stabilizer pyridostatin (PDS). Accumulation of branched intermediates, in turn, depended on the presence of the translocase SMARCAL1 and, to a lesser degree, RAD51 recombinase, indicative of fork reversal in response to prominent repeat-mediated replication fork stalling. Finally, our FRET-based studies showed that SMARCAL1 can efficiently reverse AAGGG-containing forked substrates *in vitro*, but only when the triplex-forming ability of these repeats is prevented. Altogether, we conclude that replication stalling initiated by G4-DNA formation by pathogenic (AAGGG)n repeats results in SMARCAL1-mediated fork reversal.

## METHODS AND MATERIALS

All oligonucleotides used in this study are listed in Supplementary Information.

### Plasmid Construction

Plasmids pJH5 and pJH11 were described by us previously^45^. To construct a plasmid containing the (AAGGG)_25_ repeat, primers JH9 and JH10 were first used to amplify the 434 bp-long DNA fragment from pJH5 containing the (AAGGG)_60_ repeat. This product was inserted into a 6535 bp plasmid (built upon from vector pML113) containing an SV40 origin and L-Tag gene using the NEBuilder HiFi DNA Assembly Master Mix followed by transformation into electrocompetent SURE *E. coli* cells. Ampicillin-resistant clones containing plasmids with contracted repeats were identified by repeat PCR and sequenced. A plasmid carrying the (AAGGG)_25_ repeat was subsequently used for 2DGE analysis in cultured human cells.

### Cell culturing, transfection, and DNA isolation

HEK293T cells (ATCC CRL-11268) were maintained in Dulbecco’s Modified Eagle Medium supplemented with 10% fetal bovine serum albumin and MycoZap^TM^ Plus-CL (Lonza). To study replication of the repeat-containing plasmids, 10^6^ cells were seeded in a 10 cm dish and upon reaching 50-60% confluency, were transfected with 8 µg of plasmid DNA using jetPRIME^Ò^ reagent (Polyplus). Cell culture media was refreshed on the following day. For studies with secondary structure-stabilizing compounds, pyridostatin (PDS) to 10 µM final concentration and benzoquinoquinoxaline (BQQ) to 1 µM final concentration were added to transfected cells in the dark 6 hours prior to the isolation of replication intermediates. Replication intermediates were isolated at various times post-transfection using a modified Hirt protocol. Briefly, transfected cells were spun at 340 x g for 4 min and washed with phosphate-buffered saline (PBS). Cell pellets were resuspended in 1.5 mL Hirt lysis buffer (10 mM Tris-HCl pH 7.5, 10 mM EDTA pH 8.0, 0.6% SDS) and treated with 200 µg proteinase K at 37°C for 1.5 hours. Following this treatment, NaCl concentration was increased to 1 M, and samples were left at 4°C overnight to allow for precipitation of RNA and proteins. The following day, samples were spun at 25,000 x g for 45 min at 4°C. The supernatant containing plasmid replication intermediates was treated with phenol:chloroform:isoamyl alcohol (25:24:1) followed by isopropanol precipitation and a 70% ethanol wash.

### Gene knockdown and Western blotting

For candidate gene knockdowns, at least three independent biological replicates were treated with 30 nM of target siRNA (Horizon), alongside a scramble siRNA control, 24 hours prior to and concurrently with transfection of repeat-containing plasmid. To validate gene knockdown, protein extracts were prepared upon cell detachment and PBS wash. To this end, cell pellets were resuspended in 100 µL of cold RIPA buffer (50 mM Tris-HCl (pH 7.6), 150 mM NaCl, 1% NP40, 0.5% sodium deoxycholate, 0.1% SDS, 1 mM EDTA) with protease inhibitor cocktail (1X) and agitated at 4°C for 45 min. After centrifugation at 20,000 x g for 30 min at 4°C, supernatants were transferred into freshly pre-cooled tubes. Protein concentration in cell extracts was determined using the Bradford assay, and samples were stored at -80°C. For Western blotting, 30 µg of protein extract containing 5% v/v beta-mercaptoethanol and 1X SDS sample loading buffer (Sigma-Aldrich) was boiled at 100°C for 5 min and placed on ice. Electrophoretic protein separation was done using pre-made 4-12% bis-tris acrylamide gels (Fisher Scientific) at 100 V for 2 hours. Samples were then transferred to a nitrocellulose membrane using an iBlot^TM^ 2 gel transfer device, and total loaded protein was determined using Ponceau S staining.

Following blocking for 1 hour at room temperature with 5% skim milk in TBS-T, membranes were incubated with target antibodies at appropriate dilutions overnight at 4°C. After washing with TBS-T, membranes were incubated with respective secondary antibodies for 1-to-2 hours followed by another wash. Protein concentration was visualized using chemiluminescence. Antibodies used in this study were anti-RAD51 (Calbiochem PC130, 1:1000), anti-SHPRH (abcam ab80129, 1:1500), anti-ZRANB3 (Proteintech 23111-1-AP, 1:1000), anti-SMARCAL1 (Invitrogen PA5-54181, 1:1000), anti-DDX11 (Abcam ab30017, 1:1500), anti-HLTF (Proteintech 14286-1-AP, 1:1000), and anti-RECQ1 (Milipore sigma ABC1428, 1:1500).

### 2-Dimensional gel-electrophoresis and Southern blotting

To prepare DNA fragments for analysis by 2DGE, isolated replication intermediates were digested with DpnI (New England Biolabs) to remove unreplicated bacterial plasmid DNA, together with appropriate restriction enzymes to place the repeat on the descending arm of the Yarc. BsrGI and XbaI were used to generate a 2.7 kb DNA fragment from intermediates carrying the (AAGGG)_60_ run on the lagging-strand template; BsrGI and PciI were used to generate a 2.9 kb fragment with the opposite orientation of the repeat. Restriction digestions were conducted at 37°C for 5-8 hours followed by ethanol precipitation overnight at -20° C. 2DGE was performed as previously described^48^, with minor adjustments. In brief, digested and purified fragments were first run alongside an appropriate ladder in a 0.5% agarose gel, starting at 0.25 V/cm for 20 min and then at 0.85 V/cm for 19 hours. The ladder was excised, stained in 0.3 μg/mL ethidium bromide solution, and visualized to estimate replication intermediate migration. For the second dimension, these intermediates were excised and run perpendicular to the first dimension in a 1.3% agarose gel with 0.3 μg/mL ethidium bromide at 4.23 V/cm for 10 hours at 4°C. Following depurination with 0.25 M HCl for 10 min, DNA was transferred to a positively charged nylon membrane through passive diffusion using 0.4 M NaOH as a transfer buffer. After 2-3 days, membranes were washed with 2X sodium saline citrate (SSC) and prehybridized in Church and Gilbert’s hybridization buffer (G-Biosciences), rotating overnight at 65° C. The following day, a radioactively labeled probe complementary to the analyzed fragment was synthesized and added to the membrane, again rotating overnight at 65°C. To remove any nonspecifically bound probe, the membrane was washed twice with solution one (0.1x SSC, 0.1% SDS) for 15 min and then once with solution two (2x SSC, 0.1% SDS) for 8 min. After drying on Whatman paper, the membrane was exposed to a phosphor-sensitive screen for 1- 5 days. Exposed screens were visualized using an Amersham^TM^ Typhoon^TM^ phosphor-imager. Quantification of the branched intermediates (noted as “reversed forks”) was conducted using ImageJ. Lines were drawn to encompass the branched intermediates as well as the entirety of the Y-arc. Background signal was first subtracted from each quantity, and the Y-arc then served as the proxy for all isolated replication intermediates in each individual sample to which the reversed forks, or forks localized to repeat tract, were normalized (Supplementary Figure S1A). The values for genetic controls are depicted as fold-change increase or decrease in quantity relative to wild-type.

Statistical analysis and data visualization were conducted using Prism software.

### Chromatin immunoprecipitation and qPCR

24 hours post-transfection with repeat-containing plasmid, adherent cells were detached from dishes and resuspended in 10 mL of media supplemented with 1% formaldehyde solution, gently rocking at room temperature for 10 min. Crosslinking was then quenched using 125 mM glycine, and cells were spun down at 350 x g at 4°C. Pelleted cells were resuspended in cold PBS supplemented with 1mM phenylmethylsulfonyl fluoride (PMSF), spun, and washed again with PBS-PMSF solution. Crosslinked cells were then frozen at -80° C until ready for use. In preparation for immunoprecipitation, Protein A magnetic beads (NEB) were transferred to a magnetic rack and washed three times with blocking solution (PBS supplemented with 0.5 % bovine serum albumin). Beads were then incubated with 10 µg of targeting or negative control IgG antibody, rotating overnight at 4° C. Per IP, 2 million frozen cells were resuspended in 7 mL cold lysis buffer 1 (50 mM HEPES-KOH pH 7.5, 140 mM NaCl, 1 mM EDTA, 10% glycerol, 0.5% NP-40, 0.25% Triton X-100, with 1X mini protease inhibitor cocktail) and homogenized using a tissue grinder. Following centrifugation at 700 xg for 5 min at 4°C, intact nuclei were resuspended in 10 mL cold lysis buffer 2 (10 mM Tris-HCl pH 8.0, 200 mM NaCl, 1 mM EDTA, 0.5 mM EGTA, with 1X mini protease inhibitor cocktail) and gently rocked at room temperature for 10 min. Samples were once again spun at 700 xg for 5 min at 4°C. To release chromatin, nuclei were resuspended in cold lysis buffer 3 (10 mM Tris-HCl pH 8.0, 100 mM NaCl, 1mM EDTA, 0.5 mM EGTA, 0.1% Na-deoxycholate, 0.5% N-lauroylsarcosine, 1% Triton X-100, with 1X mini protease inhibitor cocktail) and agitated at 4° C for 2 hours with regular inversion and mixing. Samples were then spun down at 20,000 xg for 10 min at 4° C and transferred to fresh tubes; sample input was taken and stored at -20° C. Antibody-bead mixtures were collected with a magnetic rack and washed again with blocking solution. Beads were resuspended with isolated chromatin, and samples were incubated overnight at 4° C. Samples were retrieved the following day and washed five times with cold RIPA buffer. De-crosslinking was performed overnight in TE solution (50 mM Tris-HCl pH 8, 10 mM EDTA, 1% SDS) with 20 µg proteinase K at 65°C. To remove RNA, samples were incubated with 5 µg RNase A at 37°C for 1 hour. NaCl was added to a final concentration of 200 mM, and DNA was purified using phenol:chloroform:isoamyl alcohol (25:24:1) followed by an isopropanol precipitation.

Enrichment of target protein on isolated DNA was validated using qRT-PCR with 10 ng DNA and iTaq Universal SYBR Green Supermix (Bio-Rad). Samples were brought to 50° C for 2 min and initial denaturation proceeded to occur at 95° C for 10 min. Through 40 cycles, samples were then brought to 95°C for 15 seconds and 60°C for 1 min. Final denaturation and elongation steps first occurred at 95°C for 15 seconds and 60°C for 1min, then at 95°C for 15 seconds and 60°C for 15 seconds.

### Protein purification

Human Flag-tagged SMARCAL1 was purified as previously described^49,50^. Recombinant human RPA was expressed using the pET-11d plasmid and purified from *E. Coli* as previously described^51^. The 6xHis-tagged human RAD52 protein was purified as described^52^. Protein purity was confirmed by SDS-PAGE, mass photometry, and stoichiometric binding assays^53^.

### Preparation of Fork DNA Substrates for *in vitro* fork reversal analysis

All DNA oligonucleotides were purchased from Integrated DNA Technologies (Coralville, IA, USA). A model replication fork containing a 30 nt lagging strand gap was prepared as described previously^49,50^. The HGQ and HDNA model forks containing either a G-quadruplex in the lagging strand gap or an H-DNA formed by the gap and parental duplex were assembled from oligonucleotides CANVAS1+ CANVAS 2+ CANVAS 3+ CANVAS 4 and CANVAS 3+ CANVAS 4+ CANVAS 5+ CANVAS 6, respectively (see **Supplemental Table 1** for sequences). In each fork, a 2 bp mismatch was present on the parental strands to prevent spontaneous branch migration. To prepare the fork reversal substrates, leading or lagging parental strands were annealed with their corresponding nascent strands in TEN buffer [10 mM Tris-HCl (pH 8.0), 1 mM EDTA, and 100mM NaCl], and both mixtures were separately heated to 95°C for 5 min in a thermocycler, then slowly cooled to 55°C. Then, equal amounts of leading and lagging DNA intermediates were mixed and slowly cooled to 4°C to complete annealing of the parental- parental region. Fully annealed model forks were purified from PAGE gel using Model 442 electro-eluter system (BIO-RAD) and stored at 4°C until use.

### *In vitro* FRET-based fork reversal assay

FRET-based analyses of DNA fork reversal by SMARCAL1 in the presence of RPA and RAD52 were carried out using a Cary Eclipse fluorescence spectrophotometer (Agilent Technologies) at 25°C in the reaction buffer containing 20 mM Tris-Acetate (pH 7.5), 100 mM KCl, 5 mM MgCl_2_, 100 μg/ml BSA, 2 mM ATP, and 2 mM DTT. Measurements were initiated with buffer only (baseline), followed by the addition of 3 nM of the respective DNA substrate, dually labeled with Cy3 and Cy5 fluorophores. The Cy3 (FRET donor) was excited at 530 nm (10 nm slit width); the emission of the acceptor Cy5 and donor Cy3 fluorophores was monitored simultaneously at 660 nm and 565 nm, respectively (10 nm slit widths). FRET was calculated as:

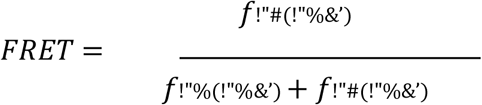

After pre-incubation with indicated amounts of RPA and/or 100 nM RAD52 for 10 min, the fork reversal reaction was initiated by addition of 1 nM SMARCAL1. Fork reversal was observed as a decrease in FRET. The FRET progress curves were fitted to a one-phase decay model using GraphPad Prism. The initial rate of fork reversal was determined as decay constant k (min^-1^) multiplied by DNA concentration (3 nM), and reversal extent was calculated as FRET span/span_max_, where span_max_ is the FRET of 100% product control (0.44 for HGQ and HDNA).

### BQQ-OP assay

BQQ-OP-mediated cleavage was performed to assess H-DNA formation in the (AAGGG)₆₀ plasmid. Plasmid DNA, 500 ng, was incubated in buffer containing 10 mM sodium cacodylate, 100 mM NaCl, and 2 mM MgCl₂. Freshly prepared BQQ-OP and CuSO₄ were mixed for 15 min at room temperature and added to the reaction at final concentrations of 1.5 µM and 2.36 µM, respectively. After incubation for 45 min at room temperature, freshly diluted 3-mercaptopropionic acid (Sigma-Aldrich/Merck) was added to a final concentration of 2 mM, and samples were incubated for 3 h at 37°C.

DNA was purified using Qiagen silica columns (QIAGEN). Untreated supercoiled plasmid, MluI-linearized plasmid, and BQQ-OP-treated plasmid followed by MluI linearization were analyzed in parallel. Samples were analyzed using a 0.7% agarose gel and visualized by SYBR Safe (Invitrogen by ThermoFisher Scientific) staining.

## RESULTS

### Pathogenic (AAGGG)_n_ repeats stall DNA replication in an orientation- and lengthdependent manner

To analyze replication of structure-prone repeats, we utilize an episome containing the SV40 origin of replication and encoding large T-antigen (L-Tag), which serves as replication initiation as well as a replicative helicase during replication elongation, and structure-prone repeats of interest^45,54^. For the current study, the (AAGGG)_60_ repeat was placed ∼1.6 kb downstream from the bidirectional replication origin, positioning the homopurine run on the lagging-strand DNA template of the clockwise replication fork (Figure 1A). We then executed 2DGE, which generates a comprehensive map of the relative populations of replication intermediates through a fragment of interest (Figure 1B)^55,56^. We’ve previously shown that replication through the (AAGGG)_60_ repeat results in an accumulation of two main populations of replication intermediates: a bulge on the Y-arc representative of forks stalled at the repeat and slower migrating intermediates corresponding to branched intermediates of the same size (Figure 1C). These branched intermediates exiting from the stall site are recognized as reversed replication forks^57,58^. Note that these intermediates migrate differently from the various recombination intermediates on the 2DGE^56,59,60^.

**Figure 1.**
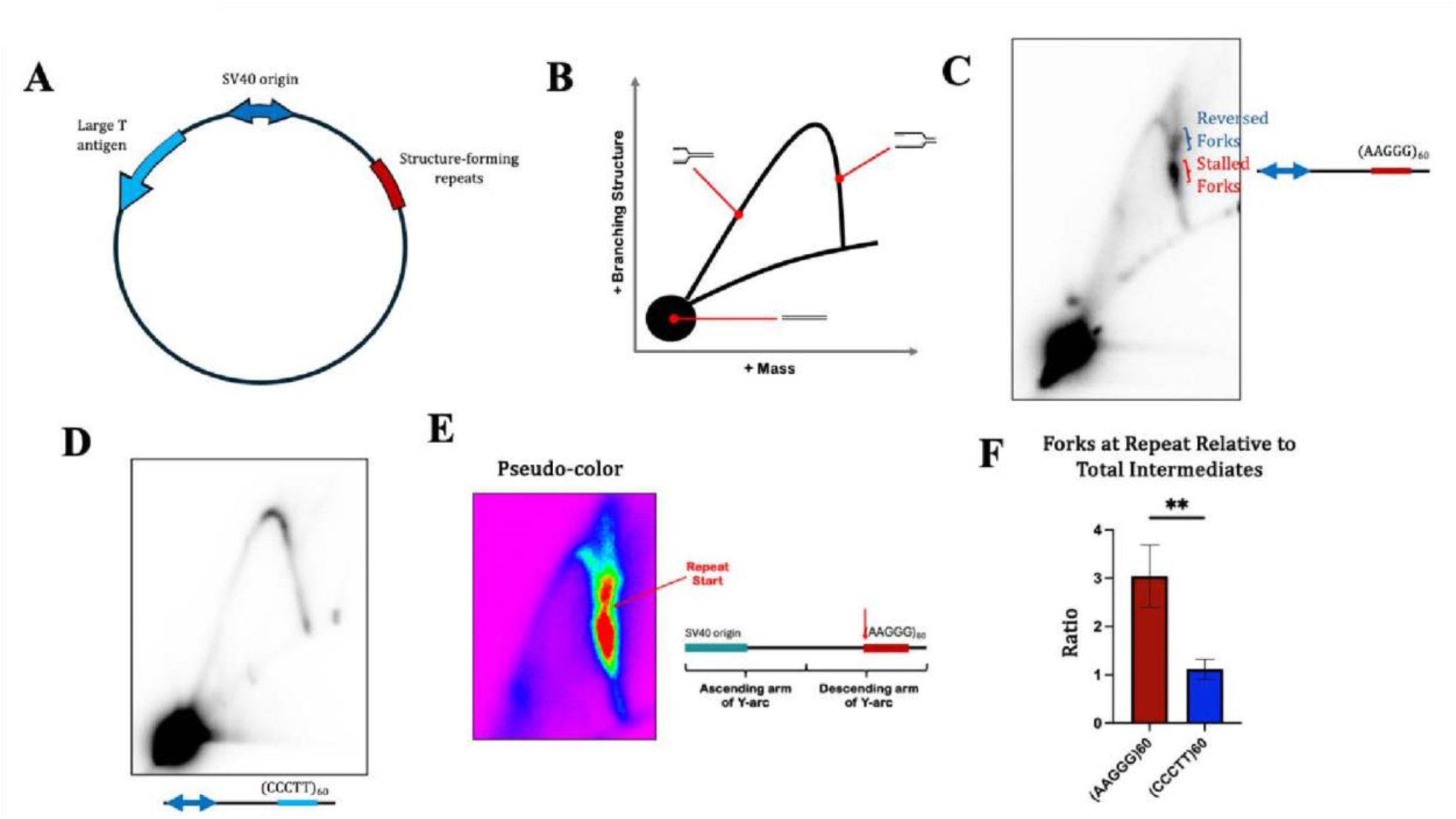
2DGE analysis to study DNA replication through AAGGG repeats. **A.** Scheme for SV40-based plasmid used in these studies. In addition to SV40 origin of replication, plasmid contains gene encoding LTag and structureforming repeats on lagging-strand template relative to rightward-moving fork. **B.** Cartoon of representative 2DGE image depicting replication forks bridging unreplicated fragments to their largely replicated counterparts. **C.** 2DGE analysis of lagging-strand synthesis of (AAGGG)_60_ repeat. **D.** 2DGE analysis of lagging-strand synthesis of (CCCTT)_60_ repeat. **E.** Pseudo- color depiction of AAGGG-mediated stalling. Note that warmer colors correspond with higher grayscale values, indicating a larger saturation of intermediates. **F.** Quantification of ratio of intermediates localized throughout repeat tract(s) relative to total intermediates isolated. Data was analyzed using an unpaired t-test with Welch’s correction, ** signifies P< 0.01.

While replication fork stalling caused by structure-prone DNA repeats was demonstrated previously^30,61–63^, fork stalling and reversal caused by the (AAGGG)_n_ repeat is significantly more severe than seen with other repeats in our experimental system^54,64^. Furthermore, replication hindrance caused by the pathogenic CANVAS repeat is strictly orientation-dependent: (CCCTT)_60_ run on the lagging-strand template instead caused no detectable replication fork stalling (Figure 1D).

To better understand where replication stalling occurs within the repetitive tract, we approximated the span of stalled intermediates based on the known length of the descending Yarc. This estimate shows that fork stalling begins at the proximal edge of the repeat and extends throughout the repeat tract (Figure 1E), which implies that either replication slowdown or recurrent restart of the replication fork occurs as it transverses through the repeat. Using this same approximation with the non-stalling orientation, forks throughout the repeat tract arise similarly to intermediates elsewhere on the analyzed fragment (Figure 1F), supporting the orientation-dependent mechanism of stalling.

To establish whether replication stalling by pathogenic CANVAS repeats depends on their length, we studied replication fork progression through a shorter (AAGGG)_25_ repeat. In this case, the intensity of forks stalling at the repeat is significantly decreased, and there are no branched intermediates above the stall site (Supplemental Figure S1B). This shows that mild fork stalling caused by the shorter repeat is not accompanied by detectable fork reversal, indicating that recovery of stalled forks occurs more easily.

### Forks stalled at (AAGGG)_n_ repeats convert into stable branched intermediates

To characterize the kinetics of replication fork stalling and reversal at the (AAGGG)_60_ repeat, we conducted 2DGE analysis of replication intermediates isolated at earlier time points of 10, 24, 30, and 36 hours post-transfection, relative to the initial 48-hour time point described above (Figure 2A). Episomal replication has barely started 10 hours post-transfection, as the Y-arc remains incomplete. At 24 hours post-transfection, episomal replication is efficient, and replication fork stalling at the repeat becomes evident; this stalling progressively intensifies at 30 and 36 hours post-transfection. Strikingly, conversion of stalled replication intermediates into branched/reversed fork intermediates occurs steadily, as they progressively accumulate posttransfection (Figure 2B).

**Figure 2.**
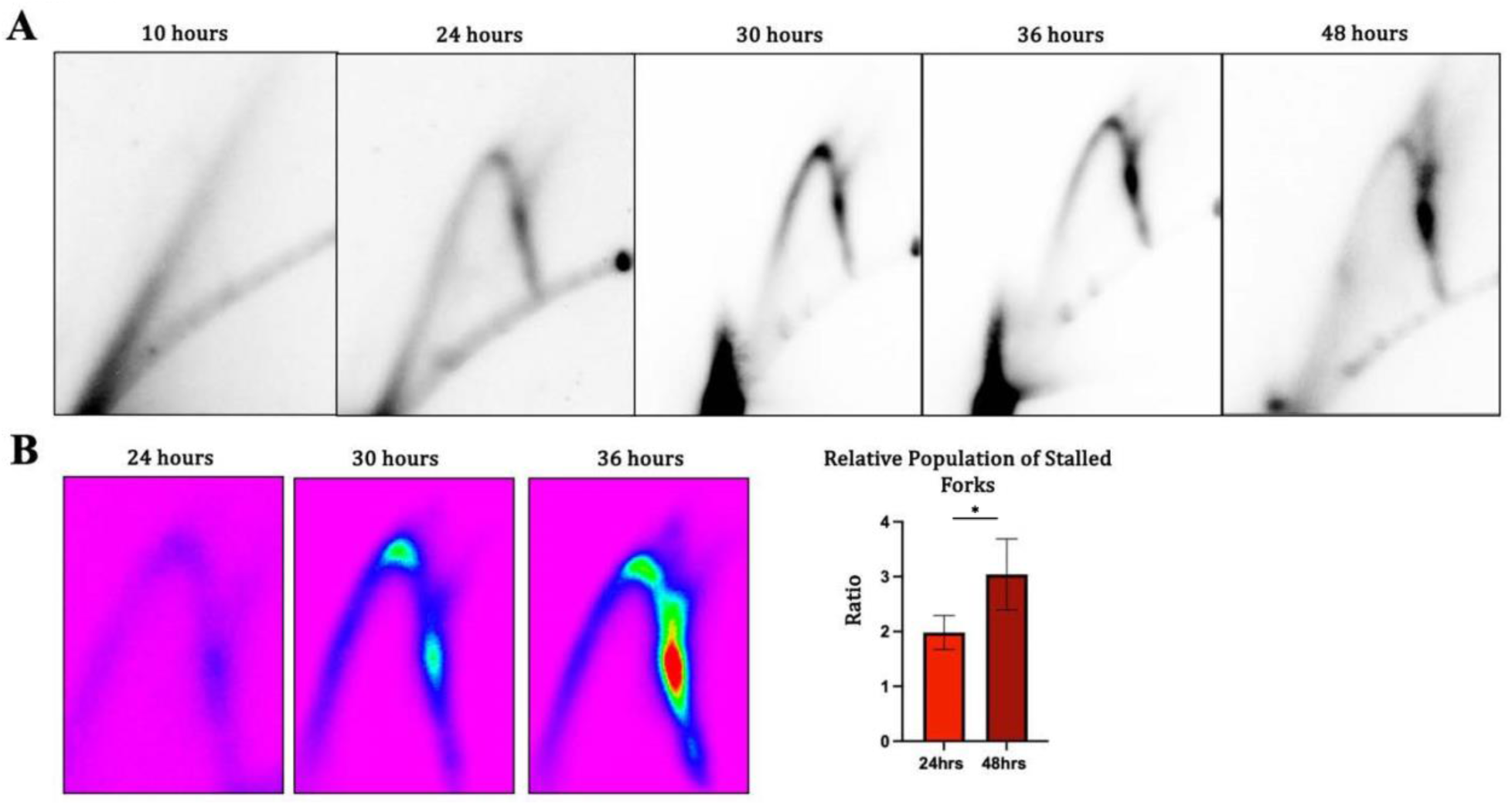
Kinetics-based assay to study intermediate formation over time. **A.** 2DGE analysis of (AAGGG)_60_ replication intermediates isolated 10-, 24, 30-, 36-, and 48-hours post-transfection. **B.** (Left) Pseudo-color depiction of focused stall sites for 24-, 30-, and 36- timepoints. (Right) Quantification of ratio of intermediates stalled at repeat tract relative to total intermediates isolated. At least three independent biological replicates were analyzed. Data was analyzed using an unpaired t-test with Welch’s correction, * signifies P< 0.05.

Consequences of replication fork stalling, particularly in response to secondary structure formation, are diverse (reviewed in depth in Sethi, A. et al. 2025)^31^, and they can differ depending on the structure of the stalled fork substrate. In our case, the (AAGGG)_n_ repeat can form an array of G4-DNAs in the lagging-strand template or triplex DNA ahead of or behind the fork. The severity of replication stalling is commonly attributed to an obstacle(s) encountered on the leading-strand template^65,66^. Note, however, that the SV40 replisome system lacks both Pol ε and the CMG helicase^67^, making it looser and likely more sensitive to the lagging-strand snags. Further, these structures may be pre-formed rather than formed de novo.

To test the latter possibility, we first used the BQQ-OP triplex-specific cleavage assay, developed in Zain’s lab, to verify whether the (AAGGG)60 repeat in our supercoiled plasmid can form a detectable triplex in vitro. In this assay, a BQQ-derived triplex-specific cleaving agent (BQQ- OP) is used. BQQ-OP recognizes H-DNA structures and cleaves double-stranded DNA at the site of triplex formation. Subsequent digestion with the MluI restriction enzyme yields two distinct DNA fragments (4186 and 8218 bps). This is exactly what we observe experimentally (Supplementary Figure S2A&B), consistent with H-DNA formation by our repeat. Altogether, we observe that the (AAGGG)_60_ repeat, being strongly prone to alternative DNA structure formation, stalls the replication fork, likely resulting in fork reversal.

### Branched intermediates arise in a SMARCAL1- and RAD51-dependent manner

It is generally believed that replication intermediates migrating directly above the stalled intermediates on the Y-arc result from fork reversal^46,57,58^. Electron microscopy confirmed that this is indeed the case for the SV40 replication fork reversal at another homopurinehomopyrimidine repeat, (GAA) ^46^. Several proteins have been shown to remodel stalled replication forks in human cells, namely the translocases ZRANB3, HLTF, FBH1 and SMARCAL1 (reviewed in Adolph M. and Cortez, D. 2024)^69^. RAD51 has also been implicated in this process by reannealing the parental duplex ahead of the fork^70^ and protecting these regressed nascent DNA arm^71^. One would expect, therefore, that inactivation of the corresponding genes will lead to a decrease in the amount of branched intermediates.

To test this idea, we analyzed the outcome of siRNA knockdown of these candidate genes on the quantity of branched replication intermediates accumulating during replication of the (AAGGG)_n_ repeat. In accord with our expectations, depletion of RAD51 and, to a greater extent, SMARCAL1 proteins resulted in a loss of branched intermediates above the stall site (Figure 3A). In the case of SMARCAL1, this loss was accompanied by a less distinct and more dispersed stall spot, suggesting slower replication progression and/or poor fork restart (Figure 3B). Note, that RAD51 is essential for accurate SV40 replication in general, and its depletion results in a loss of replication intermediates overall^72^.

**Figure 3.**
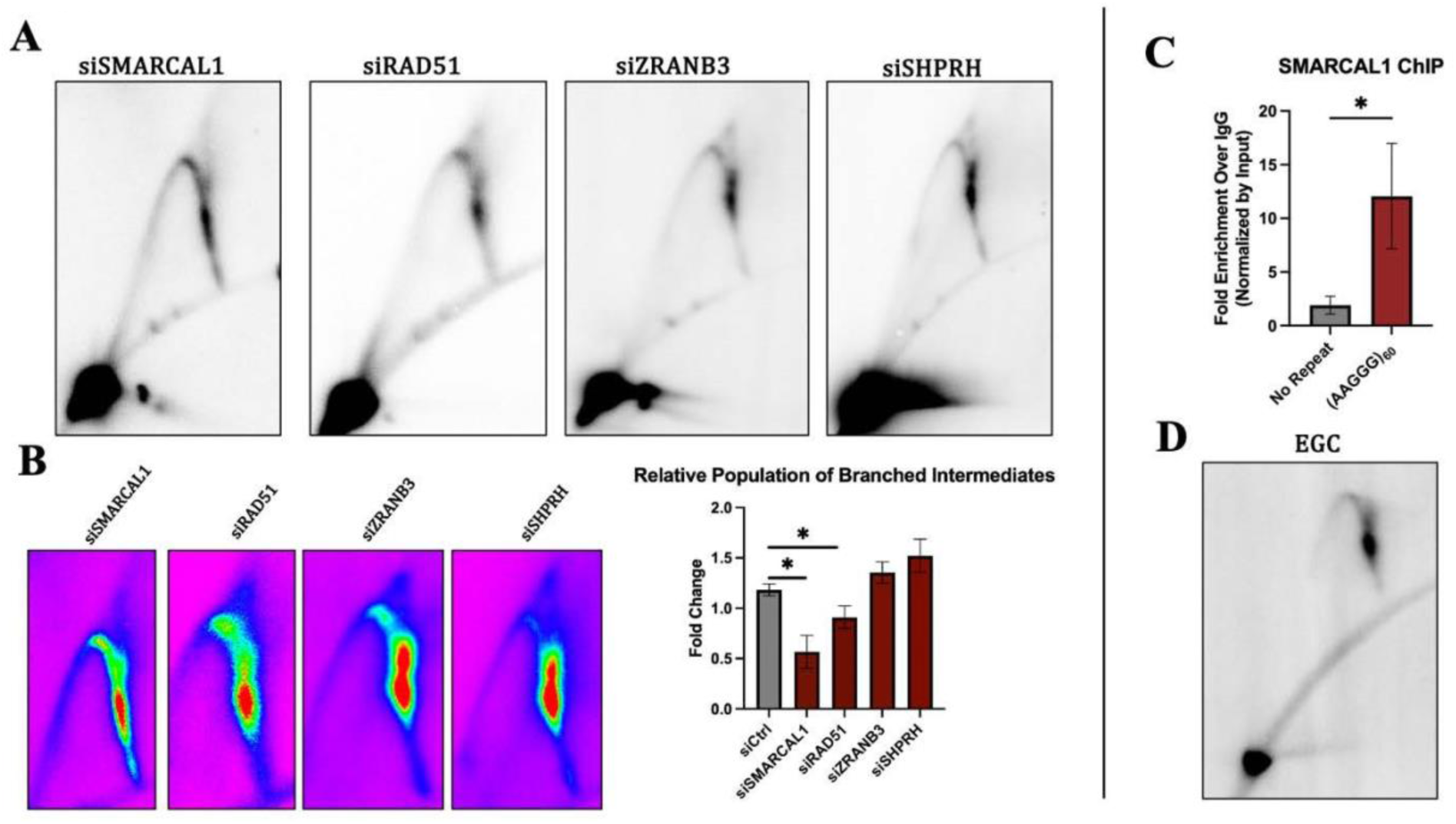
Fork restart and repair proteins involved in AAGGG replication. **A.** Genetic control 2DGE for siRNA knockdown of specific genes implicated in replication fork reversal. **B**. (Left) Pseudo-color depiction of genetic control stall sites. (Right) Relative quantifications of intermediate formation above the stall site, denoted as reversed forks, measured as fold change compared to wild type. **C**. ChIP-qPCR fold enrichment of SMARCAL1 s over non-specific IgG control between AAGGG- and no repeat-containing plasmids. **D**. Replication of (AAGGG)_60_ in presence of 5 µM RAD52 small molecule inhibitor ECG. Error bars represent standard deviation. Data was analyzed using an unpaired t-test with Welch’s correction, * signifies P< 0.05. N=3

Unexpectedly, depleting ZRANB3 did not significantly affect the fraction of branched intermediates. ZRANB3 is recruited to stalled replication forks through interaction with polyubiquitinated PCNA. The ubiquitin ligase SHPRH has been shown to ubiquitinate PCNA, functioning upstream of ZRANB3 in fork remodeling. A knockdown of SHPRH also did not reduce the fraction of branched intermediates (Figure 3A). Altogether, these findings suggest a PCNA polyubiquitination-independent pathway to fork rescue, though further experiments are required to validate this. The more diffuse stall observed with loss of SMARCAL1 (Supplemental Figure S3B) may reflect less processive translesion synthesis in the absence of a template switch pathway, though future experiments with Rad18 and other ubiquitin ligases should confirm the role of PCNA mono/polyubiquitination in this process. To validate that SMARCAL1 translocase is operating in our system, we performed ChIP-qPCR with a SMARCAL1-specific antibody. Compared to a no-repeat control plasmid, we detected roughly a 10-fold enrichment of SMARCAL1 at (AAGGG)_60_-repeat over nontargeting control IgG (Figure 3C).

RAD52 has been shown to bind stalled forks competitively both i*n vivo* and *in vitro,* thereby preventing unrestrained reversal by SMARCAL1. To determine whether RAD52 plays a similar role in our system, we used the small molecule epigallocatechin (EGC) to inhibit its activity during replication of the (AAGGG)_60_ repeat. This treatment strikingly abolished replication beyond the repeat, leaving largely only stalled forks (Figure 3D). This indicates that RAD52 likely prevents the accumulation of branched replication intermediates while helping replication throughout our episome.

Taken together, we believe that the branched intermediates accumulating above the stall site are products of RAD51/SMARCAL1-mediated replication fork reversal.

### SMARCAL1 can efficiently remodel prototypical replication forks containing G4-, but not triplex DNA

As previously mentioned, CD spectra and primer extension assays demonstrate the AAGGG repeat’s dynamic ability to form either G4 or H-DNA structures^44,45^, provided the conditions are optimal. Although it is possible that both structures could transiently form throughout duplex unwinding, SMARCAL1’s ability to initiate fork reversal at these structures may provide insight into which of them may be forming in our system. To identify what secondary structure may be formed prior to and during replication fork stalling, we evaluated the remodeling activity of SMARCAL1 *in vitro* on substrates containing the CANVAS repeat using a FRET-based assay^50,76^, and compared it to reversal of a model fork with unstructured ssDNA in the laggingstrand gap.

We constructed substrates resembling stalled replication forks with a ssDNA gap on the laggingstrand arm, which was either unstructured or contained (AAGGG)_5_ run, consistent with the replication-stalling orientation of the plasmid used for *in vivo* studies (Figure 4A&E), which we called HGQ fork. In addition, these DNA substrates contained a Cy3 dye (FRET donor) and a Cy5 dye (FRET acceptor) on the end of the leading-strand arm in the unstructured fork or at the end of the lagging-strand arm in the HGQ fork, resulting in a high FRET signal when present in the intact model fork. Upon remodeling by SMARCAL1, the FRET signal progressively decreases from ∼0.8 to ∼0.3 for the unstructured fork (Figure 4B) and from ∼0.7 to ∼0.3 for the HGQ fork (Figure 4F), making this change in signal a proxy for SMARCAL1-mediated fork reversal.

**Figure 4.**
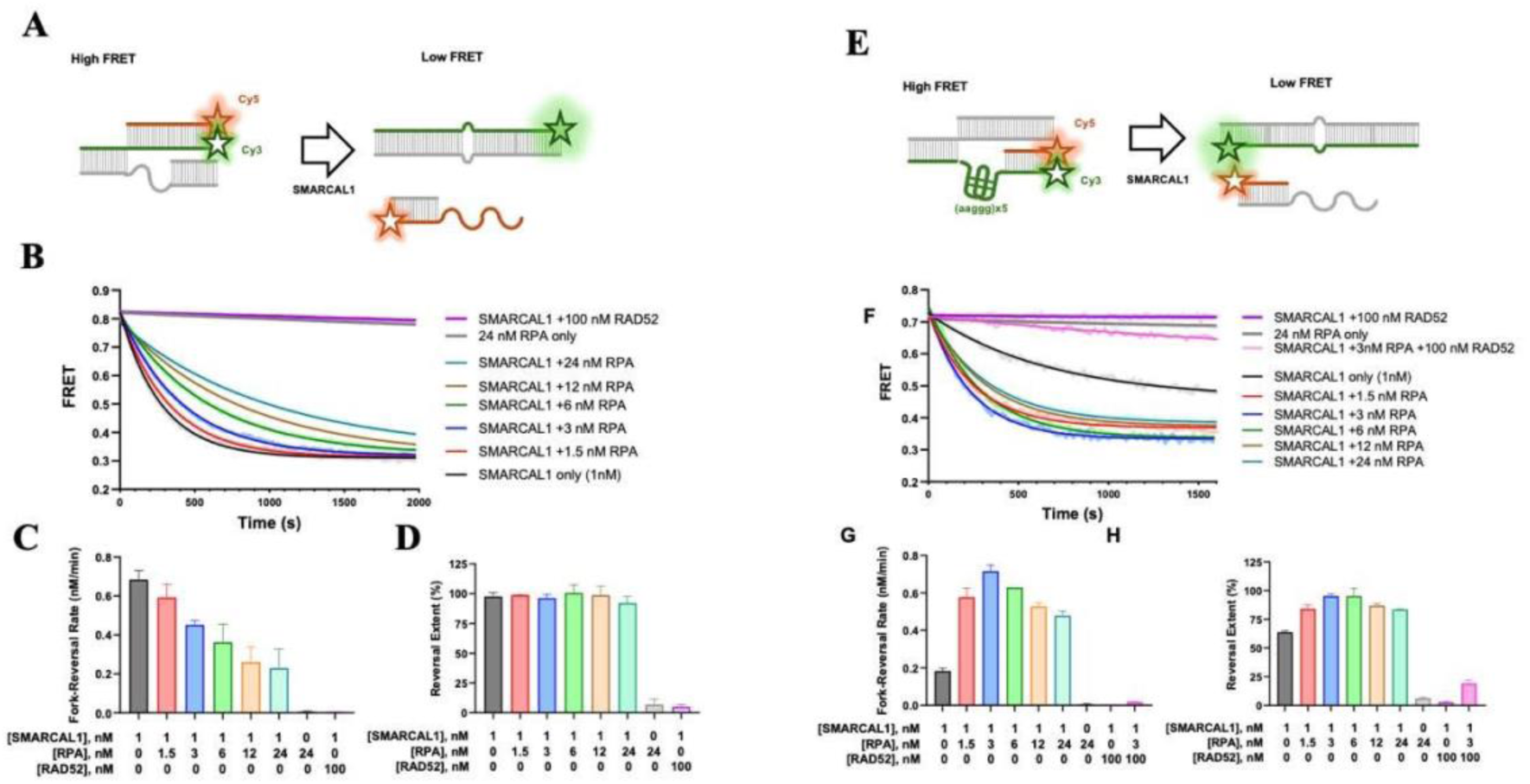
In the presence of RPA, SMARCAL1 readily remodels the model fork containing (aaggg)x5 G-quadruplex in the lagging strand gap. **A.** Model fork DNA containing unstructured lagging strand gap used in the FRET-based for reversal analysis. **B.** Representative time courses of the fork reversal (3nM) by SMARCAL1 (1nM) in the presence of indicated concentrations of RPA and RAD52. Calculated raw data are overlayed with exponential decay functions. Fork reversal time courses were fitted to single exponential functions to yield reversal rates (**C.**) and extents of the reversal reaction (D). Data in C and D are shown as average and standard deviation for two independent experiments. **E.** Model fork DNA containing five aaggg repeats in the lagging strand gap. Under our experimental condition, the sequence within the gap folds in a stable G-quadruplex. **F-H.** are the same as **B-D** but for the fork shown in **E**.

To investigate how secondary structure formation affects fork reversal, we compared two variations of the model fork with a lagging strand gap. The HGQ fork described above contained the (AAGGG)_5_ repeat in the gap region, while no repetitive DNA was present within the parental duplex, so the only potential secondary structure that can form within this fork is a G-quadruplex in the lagging strand (Figure 4E). The second HDNA fork contained an additional (AAGGG/CCCTT)_5_ repeat within the parental duplex, allowing for the substrate to fold into a triple-stranded H-DNA structure in addition to or instead of the G4 (Supplemental Figure S4A). We expect predominantly H-DNA to form in the HDNA-fork in the presence of 5 mM Mg^2+^ under ambient remodeling conditions.

In the buffer containing 100 mM KCl, SMARCAL1 (1 nM) can efficiently remodel the fork containing an unstructured lagging-strand gap; however, this activity is impacted by the presence of RPA, with higher RPA concentrations affording stronger inhibition of SMARCAL1-mediated reversal (Figure 4B-D). These observations are in line with previous *in vitro* studies, as RPA presence on lagging-strand gaps has been shown to inhibit SMARCAL1 remodeling, whereas its presence on leading-strand gaps stimulates the activity or is neutral depending on the exact model fork sequence, as the latter situation is considered more representative of fork stalling *in vivo*^50,76,78^.

The presence of the K^+^-stabilized G4 structure in the lagging strand gap of the HGQ fork slows fork reversal ∼3.5-fold and reduces the extent of the reversal reaction compared to the fork with an unstructured gap (Figure 4F-H). Low concentrations of single-stranded DNA-binding protein RPA strongly stimulate SMARCAL1 remodeling of the HGQ fork, with the fastest reversal observed at 3 nM RPA. This corresponds to a stoichiometric ratio of 1 RPA heterotrimer per 1 fork, demonstrating the importance of ssDNA in SMARCAL1’s productive engagement.

Notably, the HGQ reversal in the presence of RPA was faster than the reversal of the fork with an unstructured gap, and the inhibitory effect of higher RPA concentrations was less severe (Figure 4F-H), suggesting that forks containing G4-structures in the lagging strand template are excellent substrates for SMARCAL1-mediated reversal.

In contrast, the HDN fork appeared to be completely refractory to SMARCAL1-mediated reversal in the absence of RPA (Supplemental Figure S4B-D). While some reversal was observed in the presence of RPA, its rate was 7-fold slower than that of the HGQ fork. This likely results from sequestration of the parental duplex via the third strand of the triplex, preventing engagement of SMARCAL1 with the model fork.

Uncontrolled SMARCAL1-mediated reversal of the unstructured fork is antagonized by a DNA repair protein RAD52^50,76^ (Figure 4B-D). Similarly, RAD52 prevented SMARCAL1-mediated reversal of HGQ (Figure 4F-H) and HDNA (Supplemental Figure S4B-D) forks. These results are consistent with our in vivo EGC data described above (Fig. 3D), and they highlight the importance of RAD52 for replication fork progression past the repeat-mediated stall site.

Collectively, these data demonstrate the ability of SMARCAL1 to remodel (AAGGG)_n_ forks that form G4-DNA rather than triplex H-DNA, with effectivity dependent on RPA and structure stability.

### G4 stabilization increases repeat-mediated fork-stalling

Given that SMARCAL1 is unable to remodel triplex-forming forks *in vitro*, we hypothesized that the (AAGGG)_60_-mediated stalling observed during 2DGE analysis could be the consequence of G4-DNA formation. To test this, we turned to our previous kinetics study on intermediate formation over time (Figure 2). At 24 hours post-transfection, the dominant population of replication intermediates is stalled forks, and their concentration is not saturated to the extent observed at 48 hours (Figure 2B). Therefore, we reasoned that this 24-hour mark would be the optimal time point to identify any changes in the amount of fork stalling relative to all isolated intermediates. In our *in vitro* assays, we found that the presence of potassium, which stabilizes G4-DNA formed by the (AAGGG)_n_ repeat, decreased SMARCAL1 remodeling activity (Figure 4B). To test the effect of G4-DNA stabilization on repeat replication *in vivo*, we treated cells with the (AAGGG)_60_ plasmid with the G4-stabilizing compound PDS. At a concentration of 10 µM, treatment with PDS resulted in a significant increase in the accumulation of replication forks throughout the repeat tract (Figure 5A) relative to the control. Interestingly, despite the severity of fork stalling at the repeat, this G4 stabilization did not result in prominent fork reversal. This suggests that G4-DNA stabilization not only increases the severity of fork stalling, but also prevents SMARCAL1 remodeling of these stalled forks, in line with our FRET data (Figure 4F-H). We then treated cells with triplex-stabilizing compound BQQ, though this treatment did not result in any change to forks stalled at the repeat (Supplementary Figure S4E).

**Figure 5.**
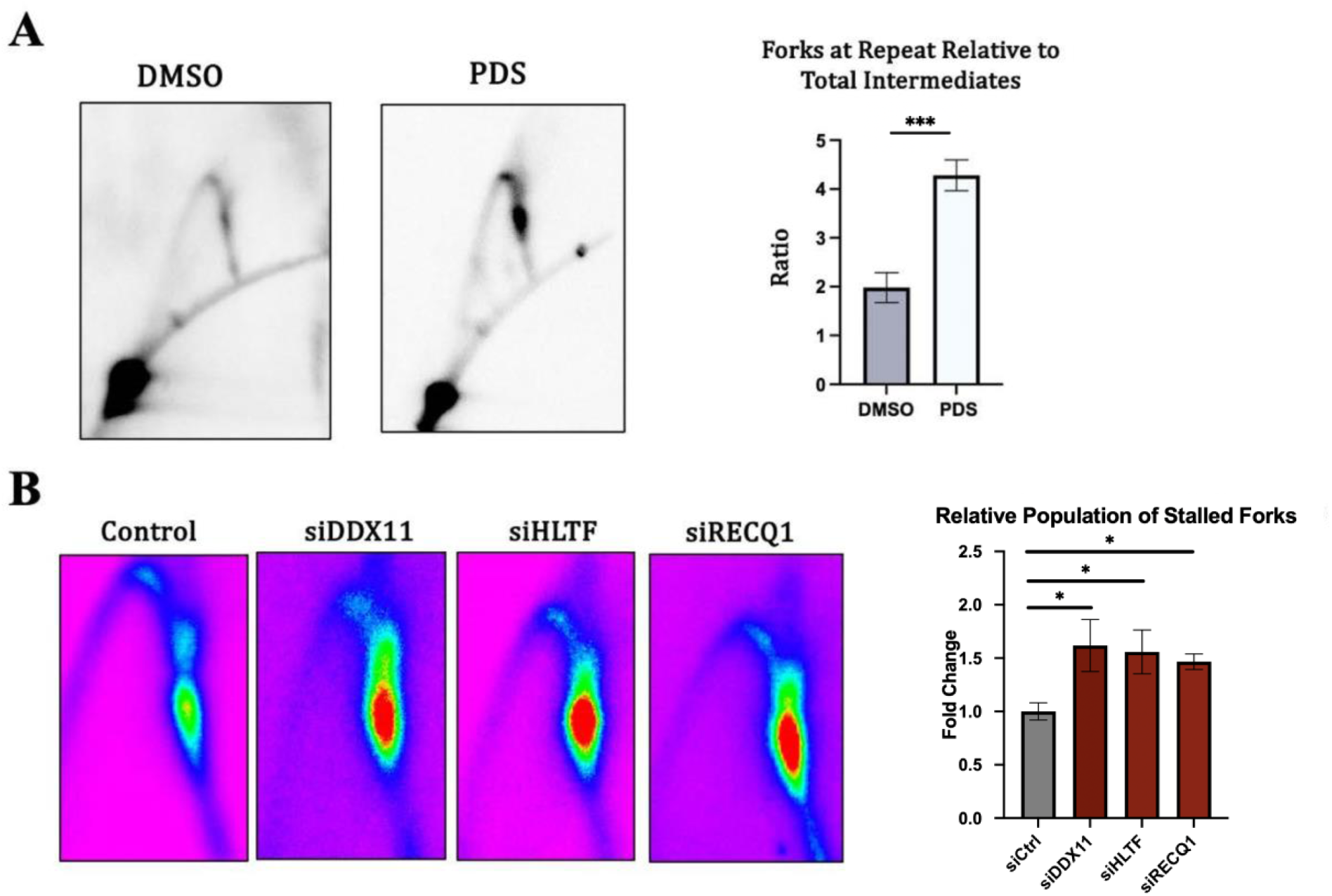
Variability of AAGGG-mediated stalling. **A.** 2DGE analysis of replication of (AAGGG)_60_ repeat in presence of G4-stabilizing compound PDS at 10 µM. Note control and PDS treated intermediates were isolated 24-hours post-transfection. **B.** (Left) Genetic control 2DGE for siRNA knockdown of accessory helicases depicted in pseudo-color. (Right) Relative quantifications of intermediate stalled throughout repeat tract, denoted as stalled forks, measured as fold change compared to wild type. At least three independent biological replicates were used for candidate gene knockdown 2DGE analysis. Data was analyzed using an unpaired t-test with Welch’s correction, * signifies P<0.05, *** signifies P< 0.001.

Provided that stalled forks are converted into stably reversed forks in untreated conditions, cells must have a means to overcome this initial obstacle. Numerous helicases have been identified as being able to unwind G4 structures, specifically in the context of replication stress in human cells^79^. These accessory helicases include but are not limited to DDX11, HLTF, and RECQ1, and *in vitro* studies have demonstrated their varying preferences for different substrates. For example, HLTF has been shown to specifically unwind G4-DNA adjacent to dsDNA^80^ but not in linear ssDNA substrates, whereas DDX11 has demonstrated the ability to resolve both G4-DNA and triplex DNA^81^. Unsurprisingly, respective knockdowns of both helicases resulted in an increase in replication fork stalling upon direct encounter with the repeat (Figure 5B). A similar phenotype can be observed from a knockdown of RECQ1. It’s important to note that in the context of secondary structure-mediated replication fork instability, RECQ1 plays a particularly dynamic role. Specifically, this helicase assists in the replication and maintenance of telomeric sequences^82^, - repetitive G-rich sequences with similar G4-forming ability as our AAGGG repeats. *In vitro*, RECQ1 has been implicated in both structure resolution and fork restart^83^, two roles indispensable for replication of sequences prone to fork stalling, like telomeric sequences^84^. Yet, more recently, several studies have also shown the importance of this helicase’s role in secondary structure resolution^85^.

## DISCUSSION

Reliable replication of structure-prone DNA sequences is essential for faithful duplication of the genome, and cellular response is complex when this process is impaired. This study sought to build on our earlier study that demonstrated replication stalling caused by the CANVAS (AAGGG)_n_ repeat^45^. Here we expand on these observations by addressing what types of replication intermediates accumulate during AAGGG-mediated fork stalling in human cells and how they arise. We have demonstrated that replication of the (AAGGG)_60_ repeat in an SV40based episome results in a dramatic replication fork stalling. Not only does this stalling persist throughout the entire repetitive tract, but it is also entirely orientation-dependent, arising only when the homopurine run is placed on the lagging-strand template. Further, the appearance of stalled forks, which becomes apparent 24 hrs post-transfection, is followed by the massive accumulation of branched replication intermediates of the same size as stalled forks, which corresponds to the electrophoretic mobility of the reversed forks. Our candidate gene analysis implies that the RAD52 protein is needed for the replication fork progression past the repeatmediated stall site, while the branched intermediates appear to be the product of SMARCAL1- and RAD51-mediated reversal of stalled forks.

PDS – a G4-DNA-stabilizing ligand – also enhances AAGGG-mediated replication fork stalling. This suggests that the formation of G4-DNA during the repeat’s replication impedes replication fork stalling. These structures are possibly resolved by accessory helicases DDX11, HLTF, and RECQ1, as indicated by a further increase in replication fork stalling upon their depletion.

To understand biochemical mechanisms responsible for the AAGGG repeat-mediated fork stalling and reversal, we created replication fork-like substrates containing the CANVAS repeat and evaluated the ability of SMARCAL1 to remodel these substrates using an *in vitro* FRETbased assay. We found that SMARCAL1 can efficiently remodel a fork-like substrate with the repeat in the quadruplex configuration, and this remodeling activity was enhanced by RPA more robustly than on the forks containing an unstructured ssDNA gap. Traditionally, SMARCAL1 has demonstrated a preference for leading-strand gaps bound by RPA as opposed to gaps on the lagging-strand where RPA can inhibit reversal by SMARCAL1^50,76,78^. These previous studies were carried out using synthetic forks with the ssDNA gap devoid of structure. In the HGQ fork, the presence of potassium stabilizes G4 structures, yet RPA can promote G4 melting and is important for SMARCAL1-mediated reversal of the HGQ fork. Surprisingly, in the presence of stoichiometric amounts of RPA, the HGQ fork appeared a preferred substrate for SMARCAL1 and was reversed faster than forks with unstructured ssDNA gaps in either the leading or lagging strand, with or without RPA. In contrast, SMARCAL1 was completely unable to remodel an HDNA fork, in which the AAGGG repeat forms an intramolecular triplex, and this remodeling shortcoming was only partially rescued by RPA in high concentrations (Supplemental Figure S4B-D). We conclude, therefore, that sequestration of the parental duplex within the triplex interferes with the engagement of SMARCAL1 on the fork.

Taken together, we hypothesize that a single-stranded (AAGGG)_n_ run that appears during lagging-strand synthesis through the repeat leads to G4-DNA formation, which in turn causes replication fork stalling (Figure 6A&B). This stall can be bypassed in the presence of the RAD52 protein and accessory replicative helicases, or it can be remodeled into a reversed fork by the SMARCAL1 and RAD51 proteins (Figure 6C). During fork reversal, the repetitive 3’ end of the nascent strand may fold back onto the repetitive duplex, forming a strong R*RY triplex that prevents both nucleolytic degradation and restoration of the reversed fork (Figure 6D). Future experiments should determine the role of fork protectors like BRCA2 in the formation and fate of reversed forks. Furthermore, given that the SV40 origin is bidirectional, stalled forks are likely rescued by the second convergent fork, explaining the absence of any 2n-sized X-spot. This may be due in part to the extreme orientation-dependent stalling of this repeat, as opposed to what we and others have observed with GAA repeats.

**Figure 6.**
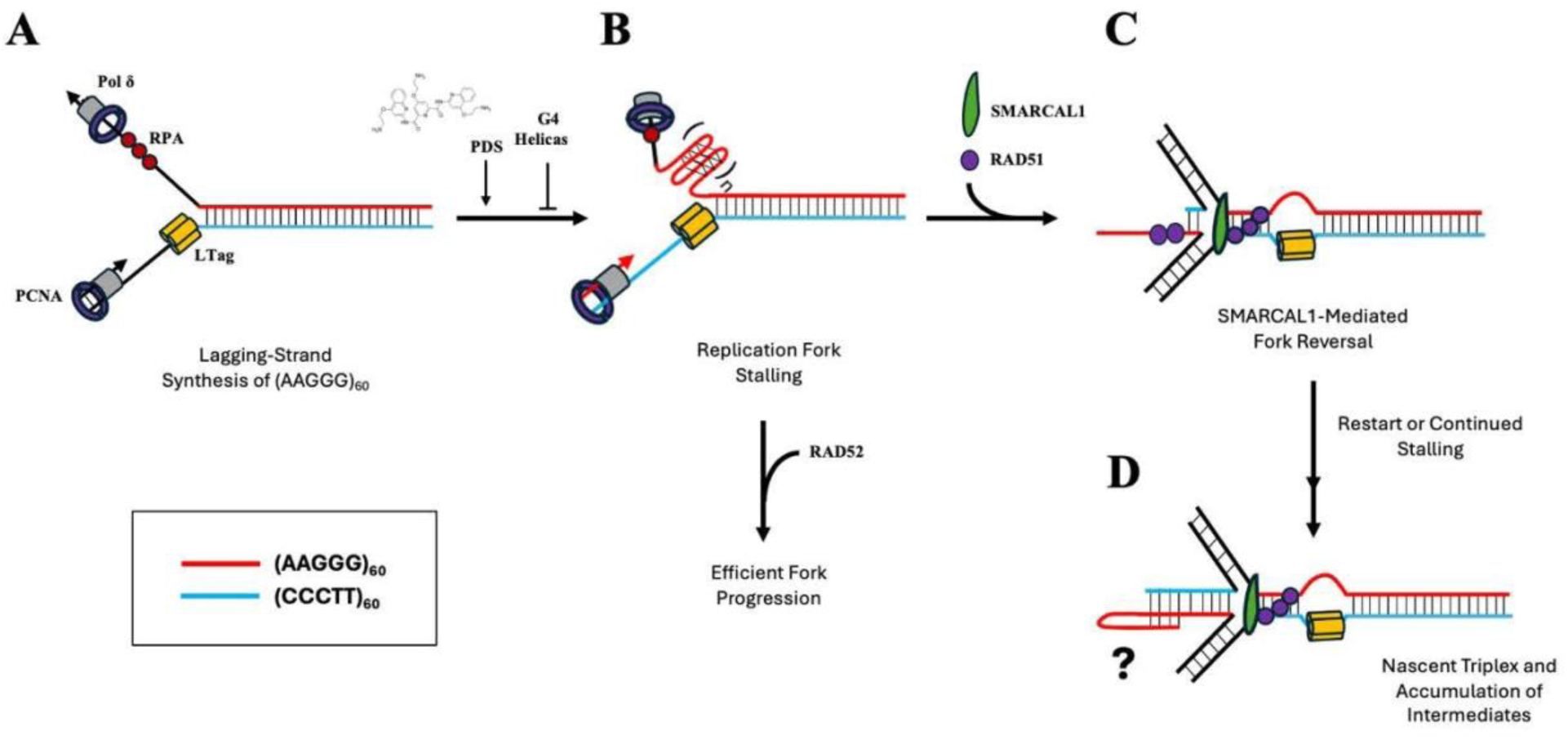

*In vitro* evidence by others^50,76,78^ suggests that SMARCAL1 is recruited by RPA to stalled forks containing unstructured single-stranded gaps in the leading-strand template. Whether and/or how the leading strand gaps are formed during the replication fork progression through the AAGGG repeat requires further studies. Our data clearly demonstrate that the stabilization of G4-DNA in the lagging strand template results in potent fork stalling followed by their reversal by SMARCAL1. One interesting possibility is that uncoupling of the replicative helicase L-Tag and DNA polymerase d in our system could create a single-stranded gap in the leading-strand template carrying (CCCTT)_n_ run, which can adopt an i-motif configuration. Supporting this idea, it was demonstrated that the leading-strand synthesis by the reconstituted replication fork *in vitro* is stalled upon i-motif formation by C-rich repeats. If this were to happen in our system, this gap might be promptly bound by the RPA^89,90^, facilitating SMARCAL1 recruitment and fork reversal. While a very fitting hypothesis, there are some important caveats to consider. Foremost, i-motifs are traditionally shown to form under acidic conditions as they require cytosine protonation, and current CD spectra do not show detectable i-motif formation by d(CCCTT)_4_ substrates under physiological conditions^91^. More recently, however, i-motifs formed by long Crich runs appeared to be stable at neutral pH^92^. Further studies are needed to address this possibility.

Exploring how DNA replication is affected by the pathogenic CANVAS repeat could shed light on paths of replication-driven genome instability, particularly when considering repeat orientation with respect to its closest replication origin. Identification of origins in human cells has greatly accelerated owing to modern technologies (reviewed in Hu and Stillman 2024)^93^.

While there is currently no strong consensus on defined origins, studies using Ok-seq data have demonstrated that initiation occurs primarily at intergenic regions adjacent to highly transcribed genes, and that replication and transcription are tightly coupled to promote codirectional progression^94,95^. To elucidate the orientation of the CANCAS repeat during replication, we analyzed available Ok-seq data focusing on replication initiation sites near the *RFC1* gene in hTERT-immortalized RPE-1 cells. Several kb upstream from the RFC1 TSS, we observed a steep conversion from predominantly Watson-mapped to Crick-mapped Okazaki fragments, indicating a replication origin (Supplemental Figure S5^95^). Not only would this align with codirectional replication and transcription, but it would also place the homopurine stretch of the CANVAS repeats on the leading-strand template, thus providing a less strenuous template for DNA replication (Figure 1D). This would explain as to why cells are able to persist with long stretches of expanded alleles, though we recognize that initiation varies across cell types and this remains purely speculative. Still, this opens the possibility for fork stalling and repeat expansions upon aberrant origin firing on the other side of the repeat, as was seen for GAA repeats in Friedreich’s ataxia iPSCs^96^. While we did not detect direct expansions of the (AAGGG)_60_ repeat in our system, we did identify an opportunity for impaired fork restart in the presence of repeatmediated stress. SMARCAL1 has been previously identified as a key player in rescue of endogenous stress at telomeres, a role not shared by ZRANB3^97^, and that this role is vital within the alternative lengthening of telomeres (ALT) pathway^98^. Given the sequence similarity of the AAGGG repeat to the G-rich strand at telomeres, it is not surprising to see SMARCAL1-driven fork remodeling in our system. This is further supported by the indispensable role of Pol δ in ALT^99^, the primary polymerase in our SV40 replisome. However, rather than aiding fork rescue, SMARCAL1 may contribute to the instability of the CANVAS repeat. While speculative, this hypothesis begs the question of how SMARCAL1 operates in individuals of a premutation or pathogenic AAGGG repeat size.

Our data in a model system indicate that even relatively short CANVAS repeats could cause replication problems. An obvious follow-up to this study is to understand whether and how these repeats affect DNA replication in iPSCs derived from CANVAS carriers and patients. 2DGE analysis is implausible in this context given the relatively lower copy of replication intermediates per an individual human gene. Exponential development of technologies aimed at genome-wide mapping of replication initiation and fork-stalling sites may circumvent this problem in the future.

## Acknowledgements

We are grateful to Mitch McVey, Catherine Freudenreich, Ralph Scully, Lindsay Julio, and Tovah Day for their invaluable suggestions and discussions, and to the current and former members of the Mirkin lab for their contributions to this project, specifically Julia Hisey, Elina Radchenko, Anastasia Rastokina, and Jorge Cebrián.

## Funding

This work was supported by the R35GM130322 from NIGMS and the NSF-BSF 2153071 to S.M.M., R35GM131704 from NIGMS and R01CA232425 from NCI to M.S.; R25 YES 1R25CA273964 from NCI and the American Cancer Society DICR INTR-24-1297273-01-DICR INTR for J.M.P.; and the Swedish Research Council, Centre for Innovative Medicine (CIMED), The Brain Foundation (Hjärnfonden), and Stockholm County for R.Z.

## Notes

### Competing Interest Statement

The authors have declared no competing interest.

### Summary of Updates

This version of the manuscript has been revised to update statistical data in Figures 2 and 5.

