## Supplemental Data for "Mechanisms of fork stalling and reversal at pathogenic CANVAS (AAGGG)_n_ repeats"

### Supplemental Table 1: Oligonucleotides used in this study

| NAME | SEQUENCE | Assembled fork name |
|---|---|---|
| **CANVAS1** (82 nt) | 5'-(Cy3)- GCTACCATGCCTGCCTCAAAAGGGGAAGGGAAGGGAAGGGAA GGGAAGGGAAGGGAAGGGAAGGGAAGGGTACTCCGCTCTA-3' | HDNA |
| **CANVAS 2** (80 nt) | 5'- TAGAGCGGAGTACCCTTCCCTTCCCTTCCCTTCCC**AA**CCCTTC CCTTCCCTTCCCTTCCCCTTTTGAGGCAGGCATGGTAGC -3' | HDNA |
| **CANVAS 3** (23 nt) | 5'- CTTTTGAGGCAGGCATGGTAGCG-(Cy5)-3' | HDNA; HGQ |
| **CANVAS 4** (47 nt) | 5'- GCTACCATGCCTGCCTCAAAAGGGGAAGGGAAGGGAAGGGAA GGGTT -3' | HDNA; HGQ |
| **CANVAS 5** (75 nt) | 5'-(Cy3)- GCTACCATGCCTGCCTCAAAAGGGGAAGGGAAGGGAAGGGAA GGGAACATTCGTCGTTGAACTGTTGAATATACC-3' | HGQ |
| **CANVAS 6** (75 nt) | 5'- GGTATATTCAACAGTTCAACGACGAATG**AA**CCCTTCCCTTCCC TTCCCTTCCCCTTTTGAGGCAGGCATGGTAGC-3' | HGQ |
| **LAG#1** (80 nt) | 5'- GGTATATTCAACAGTTCAACGACGAATGTTGGTACTCCAGTGT GTAGCATATTACGAA-CTTTTGAGGCAGGCATGGTAGC-(Cy3)-3' | Lagging strand gap fork |
| **LAG#2** (80 nt) | 5'- GCTACCATGCCTGCCTCAAAAGTTCGTAATATGCTACACACTG GAGTACCGGCATTC-GTCGTTGAACTGTTGAATATACC-3' | Lagging strand gap fork |
| **LAG#14** (22 nt) | 5'-CTTTTGAGGCAGGCATGGTAGC-3' | Lagging strand gap fork |
| **LAG#15** (50 nt) | 5'-(Cy5)- GCTACCATGCCTGCCTCAAAAGTTCGTAATATGCTACACACTG- GAGTACC-3' | Lagging strand gap fork |

the underlined 2 bp mismatch is present on the parental strands to prevent spontaneous branch migration.

### Supplemental Figure S1A

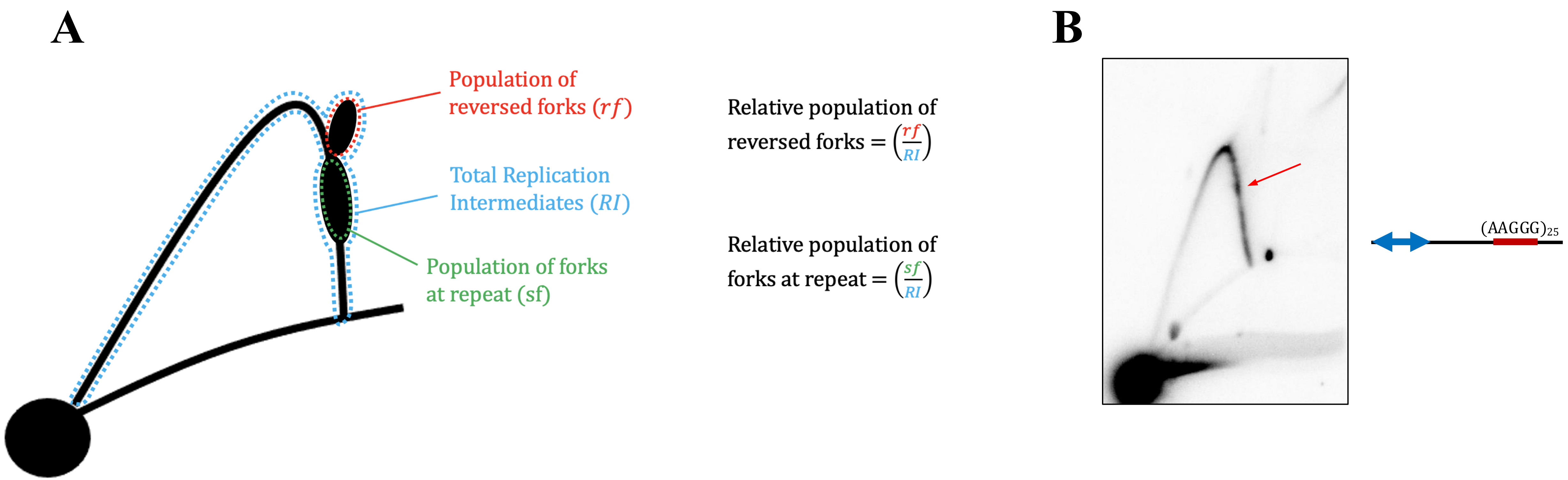


**Supplemental Figure S1. 2DGE analysis of episomal replication through CANVAS repeats.** (A) Method of quantifying the relative population of reversed and stalled forks in 2DGE. The relative population of reversed forks was approximated by taking the average density above the stall site and normalizing with respect to the total amount of replication intermediates isolated within a given sample, taken as an average of the arc density in addition to the aforementioned structured intermediates. (B) 2DGE analysis of lagging-strand synthesis of $(AAGGG)_{25}$ repeat. Arrow indicates replication fork stalling.

### Supplemental Figure S2

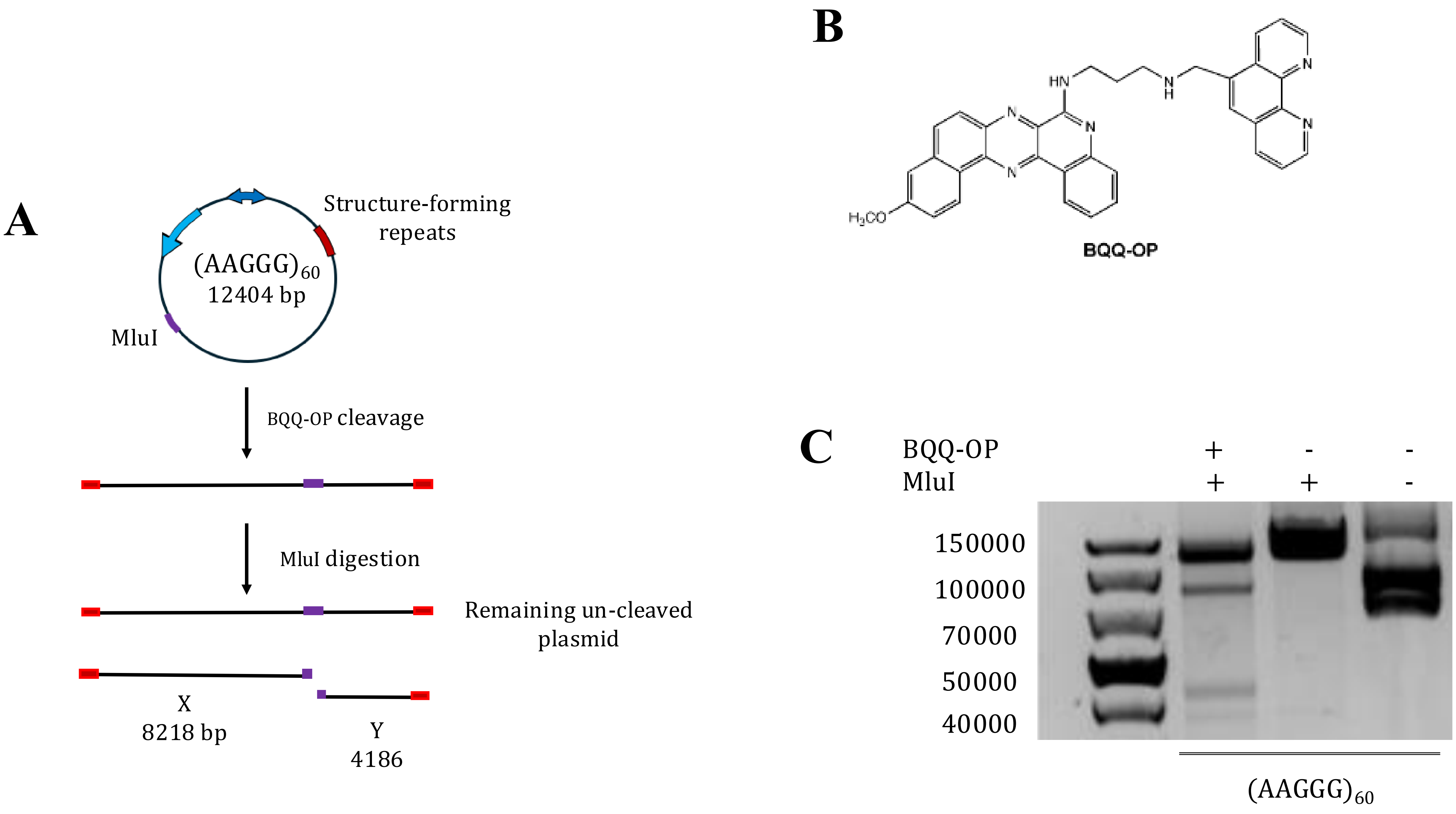

**Supplemental Figure S2. BQQ-OP-mediated cleavage of H-DNA formed by the $(AAGGG)_{60}$ repeat-containing plasmid. A.** Schematic representation of the $(AAGGG)_{60}$ plasmid and the expected DNA fragments generated following BQQ-OP-mediated cleavage at the H-DNA-forming repeat region and subsequent MluI linearization. **B.** Chemical structure of benzoquinoquinoxaline-1,10-phenanthroline (BQQ-OP). **C.** Agarose gel analysis of untreated supercoiled plasmid, MluI-linearized plasmid, and BQQ-OP-treated plasmid followed by MluI linearization. The appearance of additional DNA fragments in the BQQ-OP-treated sample indicates cleavage at the repeat-associated H-DNA structure. Data represent the percentage of cleaved plasmid DNA from three independent experiments.

**Supplemental Figure S3**

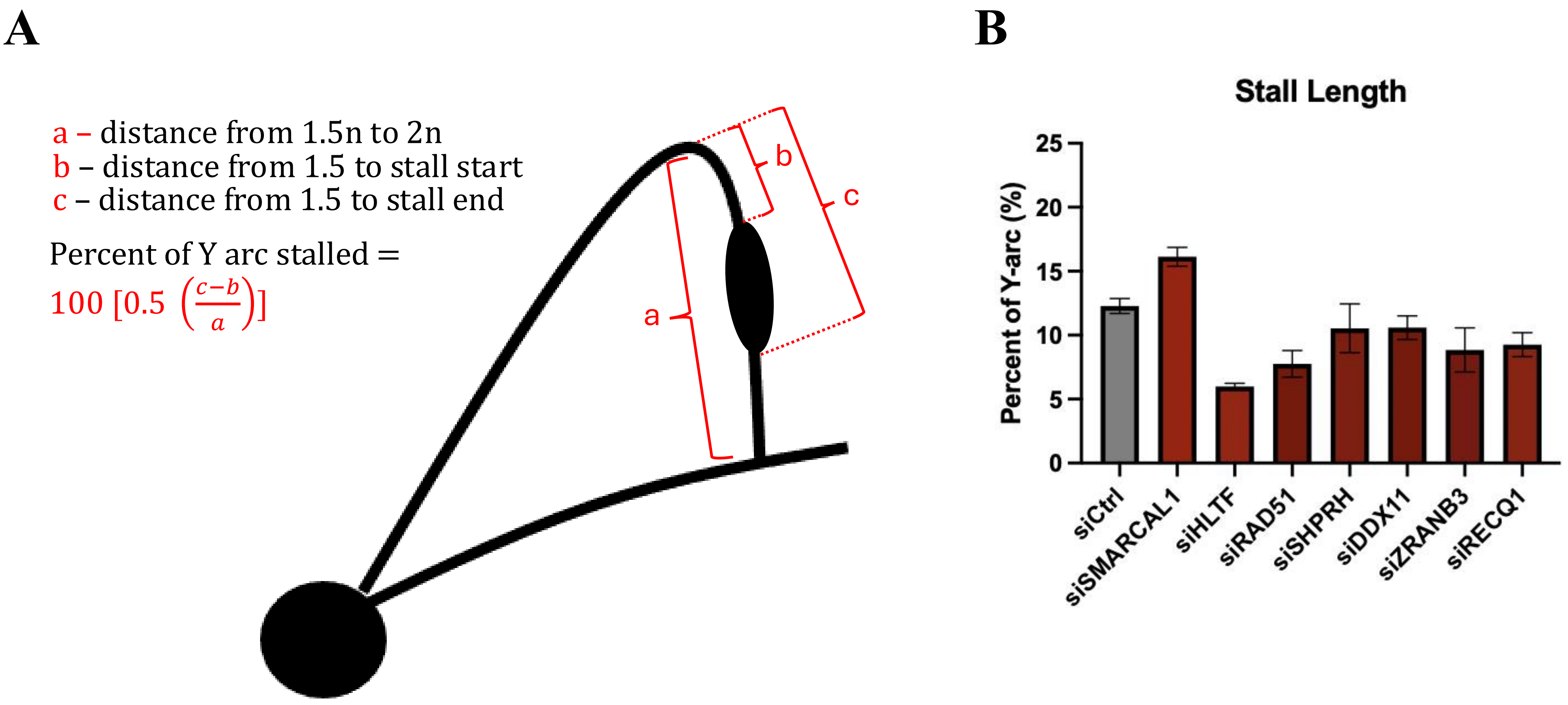


**Supplemental Figure S3. Estimation of stall extent throughout analyzed fragment. A.** Method of approximating extent of replication fork accumulation depicted as percent of Y arc stalled. Given that the descending arm is representative of replication of the latter 50% of the analyzed fragment, this provides a reference point for the relative length of fork stalling. **B.** Stall length of fork reversal and G4 accessory helicase genetic controls.

### Supplemental Figure S4

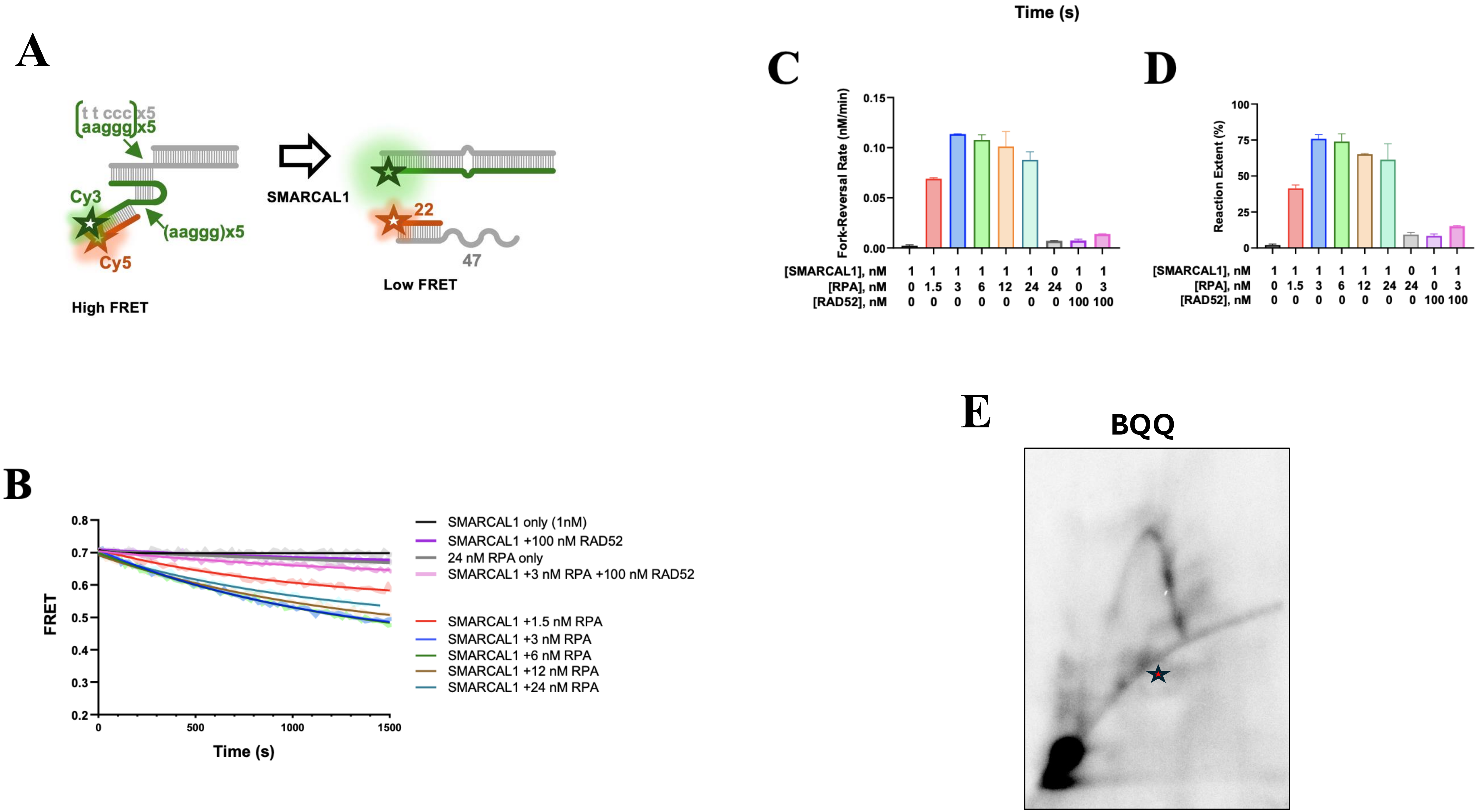

**Supplemental Figure S4. H-DNA at stalled fork is refractory to SMARCAL1-mediated reversal. A.** Model fork DNA containing five aaggg repeats in the lagging strand gap and additional five repeats in the adjacent duplex region. The repetitive DNA at the fork predominantly assumes H-DNA conformation. **B.** Representative time courses of the fork reversal (3nM) by SMARCAL1 (1nM) in the presence of indicated concentrations of RPA and RAD52. Calculated raw data are overlayed with exponential decay functions. Fork reversal time courses were fitted to single exponential functions to yield reversal rates **C**. and extents of the reversal reaction **D**. Data in C and D are shown as average and standard deviation for two independent experiments. **E.** 2DGE analysis of replication through $(AAGGG)_{60}$ upon treatment with triplex-stabilizing compound BQQ with intermediates isolated 24 hours post-transfection. Hypothesized Collapsed forks are marked with a star.

Supplemental Figure S5

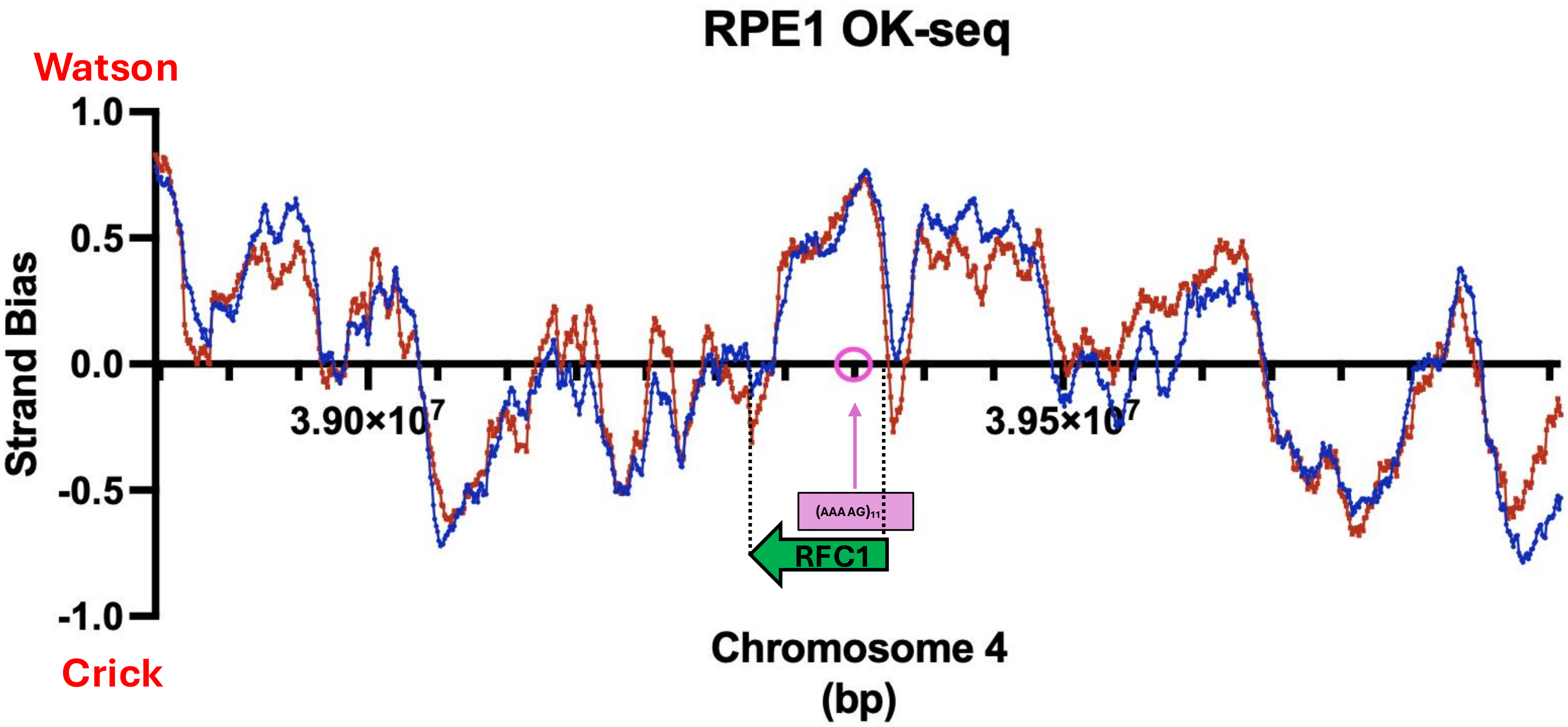


**Supplemental Figure S5. Okazaki fragment sequencing (OK-seq) data of RFC1 gene in untreated RPE1 cells.** Strand-smoothed average bias of Ok-seq data calculated from published Watson and Crick mapping of human chromosome 4 in untreated RPE1 cells (PMID: 30598550). Replicates are depicted in red and blue, respectively, with RFC1 gene and nonpathogenic $(AAAAG)_{11}$ reference allele labeled appropriated.

Supplemental Figure S6

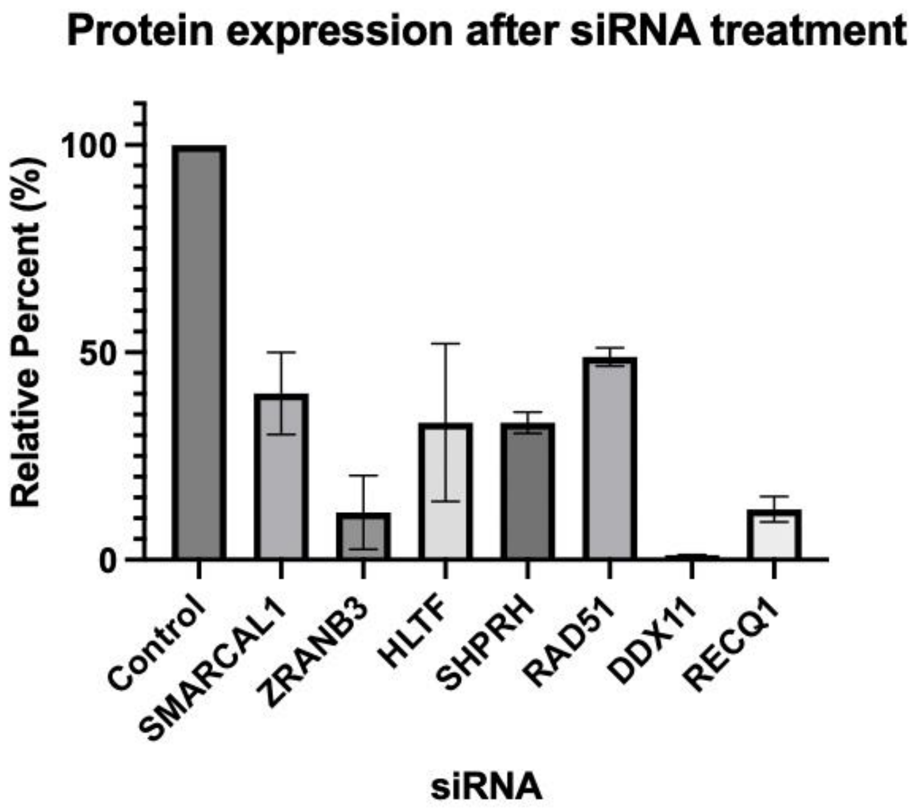


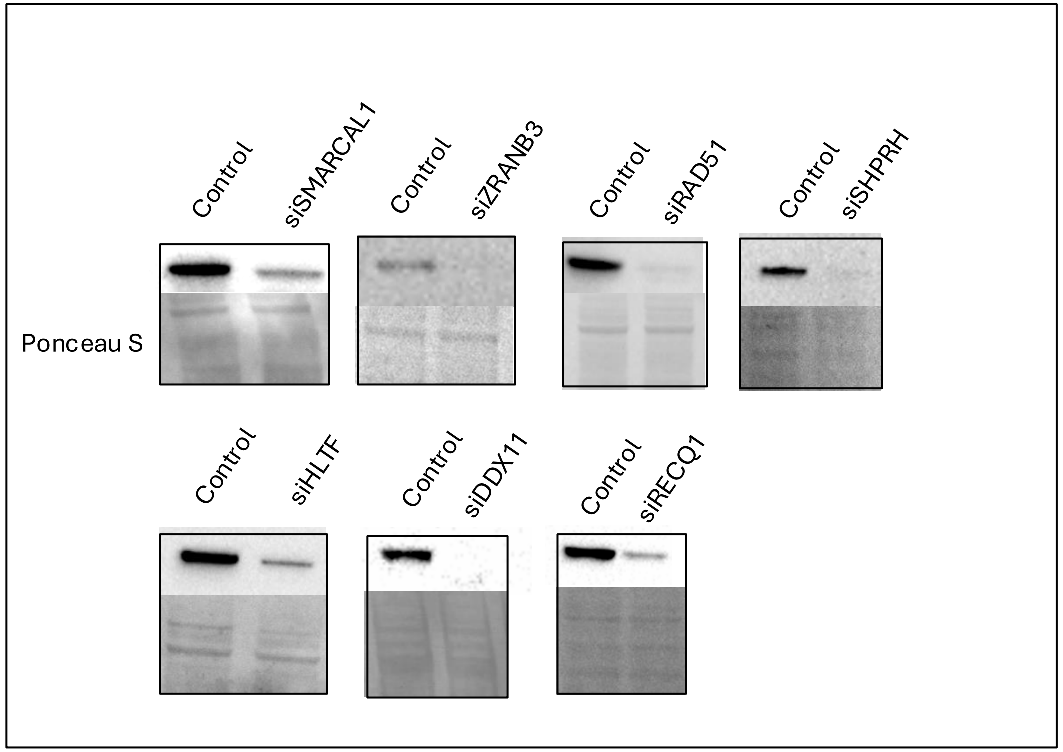


**Supplemental Figure S6. Protein expression of genetic controls.** Quantification of the relative expression (left) and western blots (right) of fork reversal-promoting and G4 helicase proteins of interest. Total protein was detected using Ponceau S staining and antibodies used in this study are listed in Methods and Materials.
